# Genome assembly and annotation of the tambaqui (*Colossoma macropomum*): an emblematic fish of the Amazon River basin

**DOI:** 10.1101/2021.09.08.459456

**Authors:** Alexandre Wagner Silva Hilsdorf, Marcela Uliano-Silva, Luiz Lehmann Coutinho, Horácio Montenegro, Vera Maria Fonseca Almeida-Val, Danillo Pinhal

**Author notes:** Equally contributed to this work (Alexandre Wagner Silva Hilsdorf, Marcela Uliano-Silva). These authors contributed equally to the work. Corresponding authors, Alexandre Wagner Silva Hilsdorf, Danillo Pinhal.

## Abstract

*Colossoma macropomum* known as “tambaqui” is the largest Characiformes fish in the Amazon River Basin and a leading species in Brazilian aquaculture and fisheries. Good quality meat and great adaptability to culture systems are some of its remarkable farming features. To support studies into the genetics and genomics of the tambaqui, we have produced the first high-quality genome for the species. We combined Illumina and PacBio sequencing technologies to generate a reference genome, assembled with 39X coverage of long reads and polished to a QV=36 with 130X coverage of short reads. The genome was assembled into 1,269 scaffolds to a total of 1,221,847,006 bases, with a scaffold N50 size of 40 Mb where 93% of all assembled bases were placed in the largest 54 scaffolds that corresponds to the diploid karyotype of the tambaqui. Furthermore, the NCBI Annotation Pipeline annotated genes, pseudogenes, and non-coding transcripts using the RefSeq database as evidence, guaranteeing a high-quality annotation. A Genome Data Viewer for the tambaqui was produced which benefits any groups interested in exploring unique genomic features of the species. The availability of a highly accurate genome assembly for tambaqui provides the foundation for novel insights about ecological and evolutionary facets and is a helpful resource for aquaculture purposes.

## INTRODUCTION

The Amazon basin harbors a massive freshwater ichthyo diversity throughout its rivers and tributaries, with 2,406 validated freshwater native fish species from 232,936 georeferenced records [1]. *Colossoma macropomum* is regarded as the largest Characiformes representative found across the Amazon River and its tributaries, with individuals reaching one meter in total length and 30 kg in weight [2] (Figure 1). This species is known by different common names, such as tambaqui in Brazil and cachama negra in Colombia. Tambaquis are omnivore/frugivore benthopelagic fish, and they have an essential ecological role as seed dispersers [3]. They are potamodromous fish, with upstream migration and reproduction taking place in the white waters along the woody shores between November and February [4]. The tambaqui is an important food and income source for Amazonian fishing communities, it is the most farmed native fish species in Brazil, with a production amount to 101,079 metric tons in 2019 [5–6].

**Figure 1.**
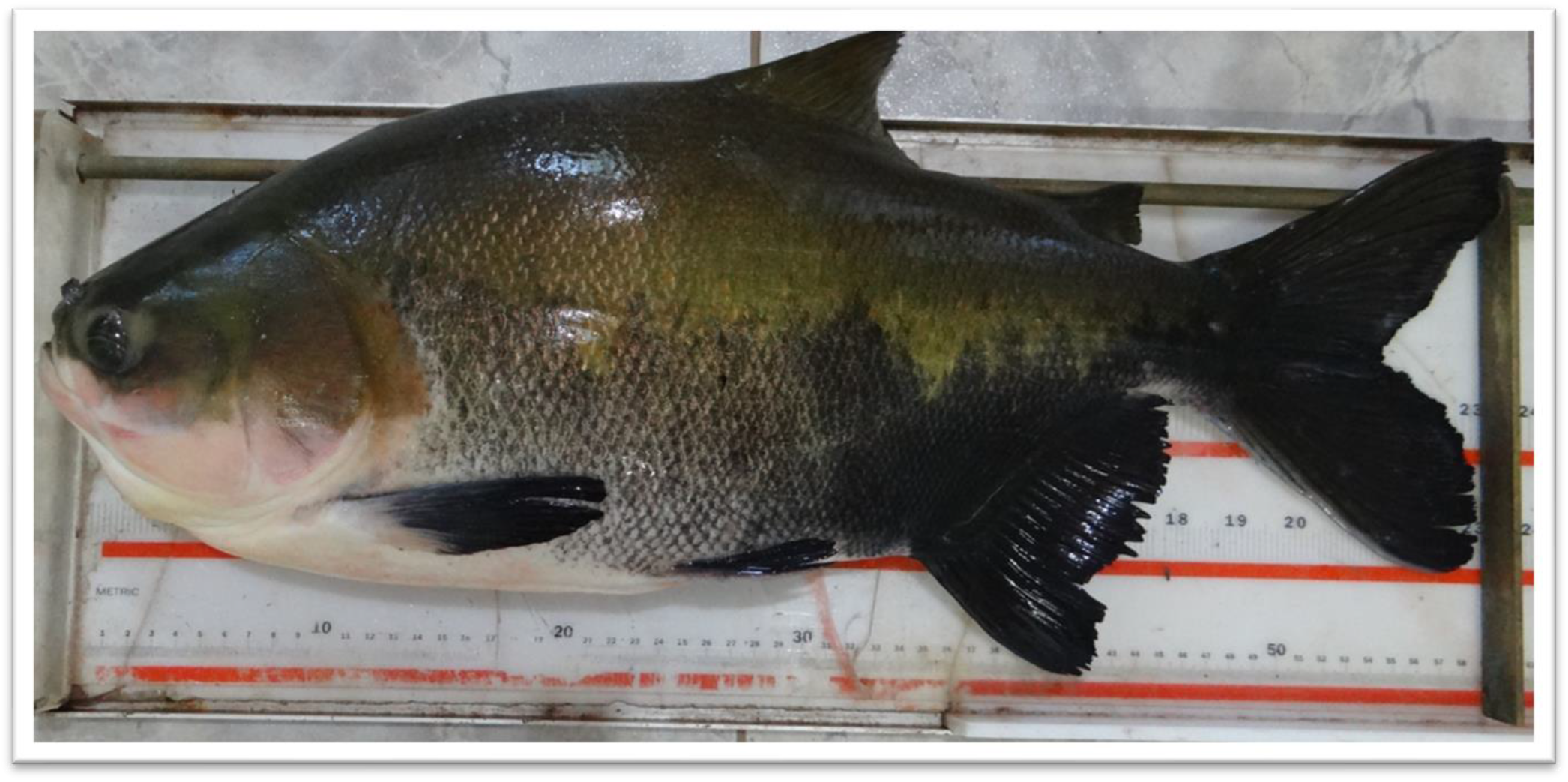
*Colossoma macropomum* individual used for the whole sequencing.

Both the key ecological and economic roles played by the tambaqui have meant that it is a comparatively well studied species, with research to date focusing on its biological adaptations to the Amazon River waters, and on the genetics of production traits to assist selective breeding programs. Transcriptomic characterization of tambaqui exposed to (i) distinct climate change scenarios and (ii) during gonadal differentiation have provided a helpful resource for the understanding of the molecular mechanisms underlying both the adaptation to a future new climate and the process of sex determination [7,8,9]. Other molecular mechanisms related to enzymatic capacity for long-chain polyunsaturated fatty acid biosynthesis have also been confirmed by a functional characterization of core genes in these processes [10,11]. Moreover, the first steps for deciphering the structure and functional dynamics of the tambaqui genome have already been taken, with large-scale SNP discovery allowing the building of a high-density genetic linkage map of the species [12], along with preliminary microRNA identification and characterization [13]. Equally pertinent are the new findings in morphology: specimens lacking intramuscular bones were identified in a fish farm in Brazil; however, the genetic and molecular mechanisms underlying the expression of such desirable phenotypes for the fish market are still unknown [14,15].

Considering the great need for increased genetic resources for the tambaqui to assist fisheries management and aquaculture [16], we present herein the first high-quality reference genome for *C. macropomum*. This complete set of DNA now represents a valuable resource for evolutionary and functional genomics studies within bony fishes, providing a window of opportunity to reveal tambaqui genome singularities and help develop molecular techniques to improve selective breeding programs.

## METHODS

### DNA isolation, taxonomy identification, and ethics statement

Genomic DNA was isolated from caudal fin-clip samples from a *C. macropomum* specimen obtained from the germplasm bank maintained by the National Center for research and conservation of freshwater aquatic biodiversity (CEPTA/IBAMA) of the Brazilian Ministry of the Environment. The specimen was a female with 3,5 Kg (Figure 1). To confirm the taxonomic status of the specimen used in this work, we have both (i) carried out an external morphological evaluation [17] and (ii) a preliminary genetic analysis of an initial Illumina run for *C. macropomum* using the kmer-matching tool Seal from BBTools package (v 37.90) [18]. We downloaded the sequences of one mitochondrial and four nuclear genes of *C. macropomum* and its two close relatives, *Piaractus brachypomus* and *P. mesopotamicus* (Supplementary Material Table S1). Then we used Seal to ascertain the number of reads with exclusive kmers matching each species’ sequences. Out of 264,813,582 reads, 1,278 matched *C. macropomum*, 62 matched *P. brachypomus* and none matched *P. mesopotamicus*, confirming the samples identification. We followed the applicable international and national ethical guidelines for the care and use of animals in research. The approval of the Ethics Committee for the Use of Animal registration is placed at the University of Mogi das Cruzes and is numbered #019/2017.

### Sequencing and assembly

Different data types were produced for the genome assembly of *C. macropomum*. High molecular weight DNA was extracted from muscle and fin clip using MagMAX CORE nucleic acid purification kit (Thermo Fisher Scientific, Carlsbad, CA, USA) to produce PacBio continuous longs reads (CLR) and Illumina paired and jumping reads (Table 2). The produced libraries were sequenced with both PacBio’s Single Molecule, Real-Time (SMRT) Sequencing technology using the Sequel system and four SMRT cells at RTL Genomics (Texas, USA) and with Illumina Hiseq2500 V4 equipment at the Functional Genomics Core Facility, Esalq-USP (São Paulo, Brazil). Illumina reads quality were checked with FastQC [19] and trimmed for adaptors and low-quality bases with BBDuk (BBBTools 37.90) (SW15-20). The genome size and heterozygosity were estimated by kmer (k=21) analysis (Figure 2A) performed with the sequenced Illumina data using meryl kmer counter, implemented in Canu assembler [20] and genome scope [21].

**Figure 2.**
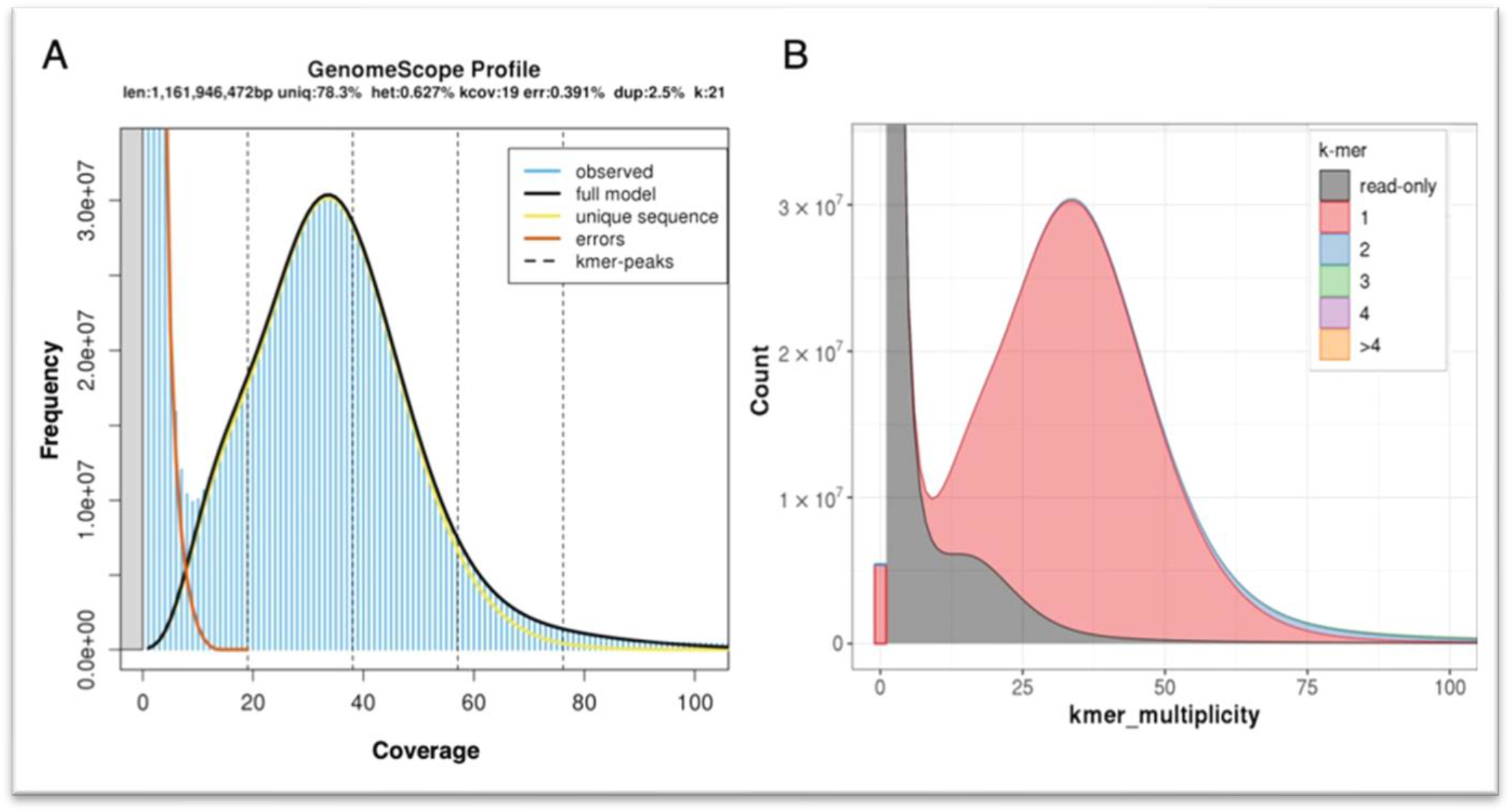
**(A)** Kmer composition of sequenced short Illumina reads (Table 1) of the tambaqui *C. macropomum.* **(B)** A merqury kmer analysis of the final tambaqui genome bases against its sequenced Illumina reads.

The 21-mers distribution of the Illumina data obeyed the theoretical Poisson distribution (Figure 2A). The genome size was estimated in 1,16 Gb with heterozygosity of 0.62%. Based on these estimations, we sequenced a 39X coverage of the tambaqui genome in long PacBio reads, and 130X in short Illumina reads (Table 1). For the genome assembly, PacBio reads were input to the assembler Flye (v2.5) [22] with parameters ‘genome-size 1.5g - pacbio-raw’. Then, the assembly was polished using the Illumina reads with the software Pilon [23] and parameters ‘frags’ for paired reads and ‘jumps’ for mate-pair reads. Finally, the assembly of the tambaqui had one round of purging with PurgeDups [24]. Purging was performed to remove any sequences representing duplicated portions of a chromosome that are erroneously kept in assemblies when the divergence level of those regions in both haplotypes is high. This has removed 1,167 contigs and 26 Mb of haplotypic retention. The final tambaqui genome was assembled into 1,269 scaffolds with a scaff N50=40Mb and a total assembly length of 1,221,847,006 bp (Table 2). A fraction of 93% of the genome is assembled on 54 scaffolds that represent the main tambaqui karyotype [25]. We have also identified the mitochondrial genome (Figure 3) within our assembled genome: it is represented by scaffold NW_023495502.1 that is 16,715 bp in length and has a conserved gene content and synteny with *C. macropomum* mitogenome available on NCBI (KP188830.1).

**Figure 3.**
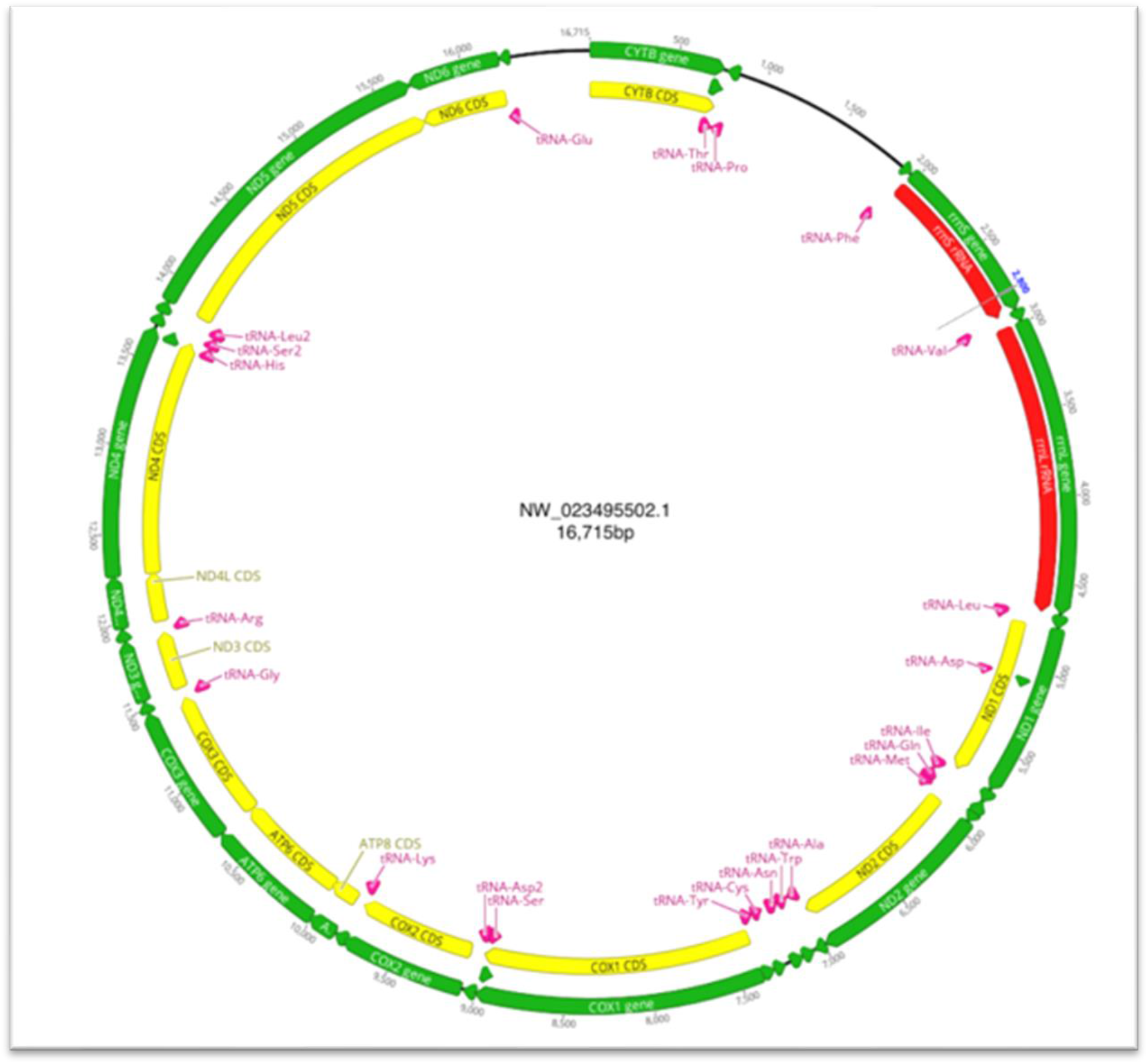
Mitogenome of *C. macropomum*

**Table 1:** Summary of genome sequencing data generated with multiple sequencing technologies. Sequencing coverage was based on the estimated genome size (1,16Gb) generated for *C. macropomum* by kmer analysis (k=21) of the Illumina sequencing data.

| Library Type | Insert<br>Size<br>(bp) | Raw<br>Data<br>(Gb) | Clean<br>Data (Gb) | Average<br>Read Length<br>(bp) | N50<br>Read Length (bp) | Clean data<br>sequencing<br>coverage (X) |
| --- | --- | --- | --- | --- | --- | --- |
| Illumina<br>(R1 and R2) | 400 | 59.08 | 52.93 | 100 | -- | 44.89 |
| Illumina<br>(R1 and R2) | 4000 | 78.81 | 57.69 | 81 | -- | 49.7 |
| Illumina<br>(R1 and R2) | 8000 | 55.59 | 41.31 | 83 | -- | 35.6 |
| Pacbio CLR | -- | 45.58 | --- | 9749 | 17667 | 39.2 |
| Total |  |  |  |  |  | 169.39 |

**Table 2:** Final statistics for the genome assembly of *C. macropomum.*

|  | Contig length (bp) | Scaffold length (bp) | Number of Contigs | Number of Scaffolds |
| --- | --- | --- | --- | --- |
| Total | 1,221,809,066 | 1,221,847,006 | 1687 | 1269 |
| Max | 26,165,397 | 63,817,184 | --- | --- |
| N50 | 5,645,235 | 40,163,545 | 54 | 14 |
| N90 | 655,952 | 2,856,822 | 300 | 33 |

### Repeat sequences and gene annotation

We identified repeat sequences in *C. macropomum* using homology-based, and *de novo* approaches. A *de novo* library of repeats was created for the tambaqui using RepeatModeler2 package [26]. This library was then combined with RepBase [27] (release 26.04), forming the final ‘teleost’ library with which *C. macropomum* genome repeats were searched. Table 3 presents the repeat summary of *C. macropomum:* 52.49% of the genome is composed of repeats, of which 49.78% are interspersed repeats. *C. macropomum* genome was submitted to NCBI for annotation. The robust NCBI Eukaryotic Annotation Pipeline uses homology-based and *ab initio* gene predictions to annotate genes (including protein-coding and non-coding as lncRNAs, snRNAs), pseudo-genes, transcripts, and proteins. Details of the pipeline are described in the NCBI Annotation HandBook (https://www.ncbi.nlm.nih.gov/genbank/eukaryotic_genome_submission_annotation/). Briefly: first, repeats are masked with RepeatMasker [28] and Window Masker [29]. Subsequently, transcripts, proteins, and RNA-Seq from the NCBI database are aligned to the genome with Splign [30] and ProSplign (https://www.ncbi.nlm.nih.gov/sutils/static/prosplign/prosplign.html). Those alignments are submitted to Gnomon [31] for gene prediction. Gnomon (i) merges non-conflicting alignments into putative models, then (ii) extends predictions missing a start and a stop codon or internal exon(s) using an HMM-model algorithm. Finally, Gnomon (ii) builds pure *ab initio* predictions where it finds open reading frames of sufficient length but with no supporting alignment detected. Models built on RefSeq transcript alignments are given preference over overlapping Gnomon models with the same splice pattern. Table 4 presents a summary of the annotation of *C. macropomum*. A detailed description of the tambaqui genome annotation can be found on the NCBI Eukaryotic Annotation Page (https://www.ncbi.nlm.nih.gov/genome/annotation_euk/Colossoma_macropomum/100/).

**Table 3.**
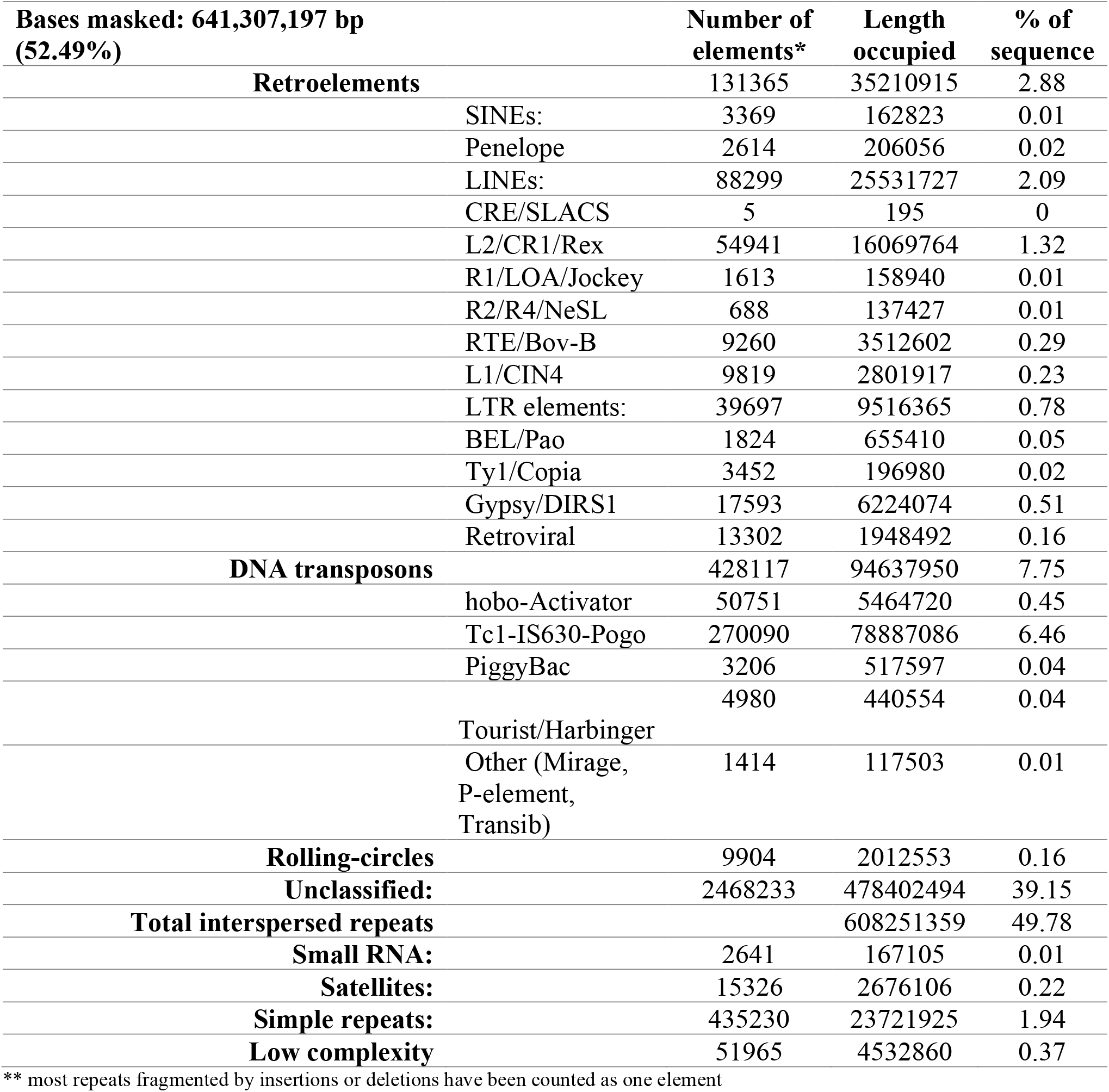
Repeat annotation: Annotation of repeats done for *C. macropomum* with a *de novo* library built with RepeatModeler added to a Repbase teleost library. The final library was used as input to RepeatMasker.

**Table 4.** Summary of the annotated features of *C. macromapum* genome

| Feature | *Colossoma macropomum* |
| --- | --- |
| Genes and pseudogenes | 31,149 |
| protein-coding | 26,670 |
| non-coding | 3,279 |
| CDSs |  |
| fully-supported | 43,938 |
| With >5% ab initio | 1,648 |
| partial | 267 |
| Protein assigned RefSeq(XP_) | 43,618 |
| Mean CDS length (bp) | 2,011 |
| Mean intron length (bp) | 2,631 |
| Mean exon length (bp) | 280 |
| Mean exon per gene | 12.02 |

Detailed annotation report can be found at:
https://www.ncbi.nlm.nih.gov/genome/annotation_euk/Colossoma_macropomum/100/#BuildInfo

## RESULTS AND DISCUSSION

All sequencing data is available on NCBI under the BioProject PRJNA702552, via SRA accession numbers SRX10122091 to SRX10122101. The assembled genome and sequence annotations are available on NCBI with the accession number GCF_904425465.1. The genome sequence and the annotation files - including CDS and proteins - can be downloaded from the NCBI FTP server (https://ftp.ncbi.nlm.nih.gov/genomes/all/GCF/904/425/465/GCF_904425465.1_Colossoma_macropomum/). Finally, a genome DataViewer was built for the tambaqui and can be accessed at https://www.ncbi.nlm.nih.gov/genome/gdv/browser/genome/?id=GCF_904425465.1. This browser is ideal for further exploration of the tambaqui genome especially from groups that are not specialist bioinformaticians, such as geneticists working on selective breeding programs.

### Evaluating the completeness of the genome assembly and annotation

The final assembly of the tambaqui is 1.2 Gb with a scaffold N50 size of 40.163 Mb (Table 2). Figure 2A shows the DNA kmer prediction of genome size done with the Illumina reads produced to polish this assembly. Further, Figure 2B presents a merqury [32] kmer plot of the final assembly: merqury produces a mapping-free evaluation of kmer completeness in genomes by comparing the assembly kmers with raw reads for the specimen. In this case, we used the high-quality Illumina reads (Table 1) to plot the merqury evaluation against the genome kmers. Figure 2B shows that (i) the kmers in the genome are in accordance with its Illumina read kmers, (ii) the assembly kmers have the same distribution of the raw reads kmer (2A), and that (iii) most of the assembly kmers (pink color) are unique in the genome, showing that the final assembly of the tambaqui has low levels of haplotypic retention (blue color). The final phred-like merqury QV score is 36.73 (QV=36.73), meaning that the tambaqui assembled bases are more than 99.9% accurate. The merqury completeness score shows that 89.31% of kmers in the Illumina reads are present in the assembly, which is a good recovery of kmers for a species with 0.6% heterozygosity.

For the tambaqui genome, 93% of the assembled bases are present in the largest 54 scaffolds. We have performed a first nucleotide synteny analysis of BUSCO genes found in the first 54 scaffolds of *C. macropomum* against the BUSCO genes on genome of *Ictalurus punctatus* [33] using busco2fasta (https://github.com/lstevens17/busco2fasta) and Circos [34]. The synteny is presented in Figure 4. *C. macropomum* and *I. punctatus* shared a common ancestor ~150 million years ago [35]. The image shows a good degree of synteny in terms of BUSCO genes, for a number of times entire chromosomes are syntenic. Supplementary Figures S1 and S2 show similar analysis with *C auratus* [36] and *Astyanax mexicanus* [37] of different levels of relatedness to *C. macropomum* demonstrating the potential of this highly contiguous genome for studies of chromosome evolution.

**Figure 4.**
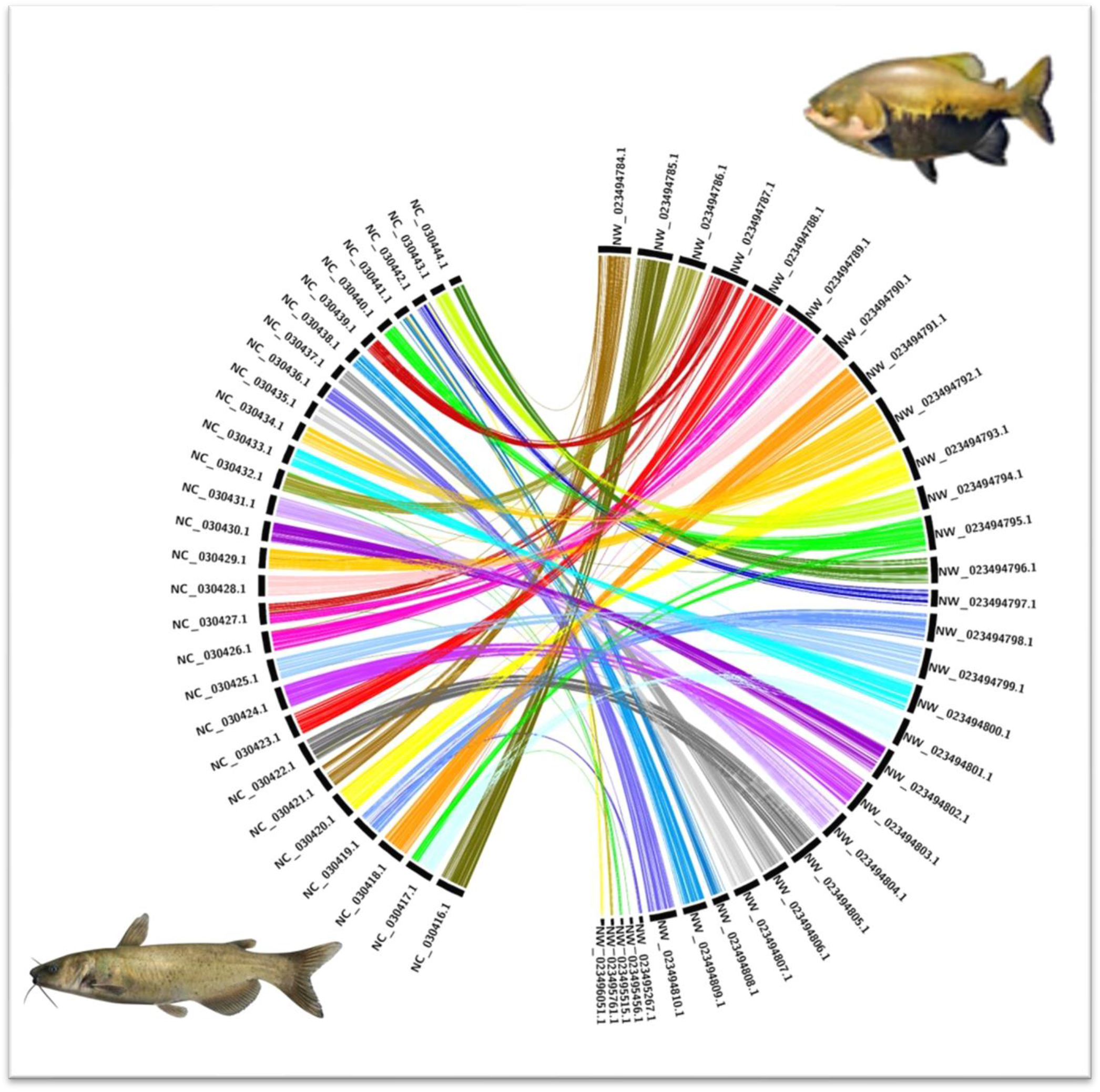
BUSCO genes synteny of *C. macropomum* (tambaqui; on the right side) and *Ictalurus punctatus* (channel catfish; on the left side). Synteny analysis of single copy genes reveal low conservation of homologous gene order between the species. The majority of *C. macropomum* genes are pulverized into several linkage groups of *I. punctatus* genome, which may reflect different genome evolutionary events experienced by them.

Finally, we have performed a general gene presence analysis of *C. macropomum* genome using the BUSCO software [38] (v5.0.0) and the OrthoDB [39] library actinopterygii_odb10. BUSCOv5 has recovered 96.5% of complete BUSCO genes out of 3,640 genes searched, where 95.5% were complete and single copy, 1.0% were duplicated, 1.0% were fragmented, and 2.5% were missing - once more demonstrating the quality of the tambaqui assembly

### Gene family identification and phylogenetic analysis of *C*. *macropomum*

To identify gene families among *C. macropomum* and other species, we downloaded the whole genome predicted protein sequences from the NCBI Eukaryotic Annotation Page of *Oreochromis niloticus* (GCF_001858045.2), *Carassius auratus* (GCF_003368295.1), *Danio rerio* (GCF_000002035.6), *Lates calcarifer* (GCF_001640805.1), *Cyprinus carpio* (GCF_000951615.1), *Rhincodon typus* (GCF_001642345.1), *Poecilia formosa* (GCF_000485575.1), *Ictalurus punctatus* (GCF_001660625.1), *Astyanax mexicanus* (GCF_000372685.2), *Oncorhynchus mykiss* (GCF_013265735.2) and *Pygocentrus nattereri* (GCF_001682695.1). We then input this data to Orthofinder [40] (v2.5.2). From all of the proteins imputed from the 12 species, Orthofinder has assigned 97.3% of the proteins to 31,794 orthogroups. There were 10,939 orthogroups with all the species present, and 33 of them consisted of single-copy genes. Those 33 single-copy orthologs were used to generate a phylogeny (Figure 5). First, the single-copy were aligned with MAFFT [41] (v7.455), and alignments were trimmed with trimAL [42] (v1.4. rev15). Then, a supermatrix was created using geneStitcher.py [43], which was imputed to PhyML [44] for a phylogeny with the amino acid substitution model LG and 100 bootstrap replicates. The phylogeny presented herein (Figure 5) is consistent with other studies [45–46].

**Figure 5.**
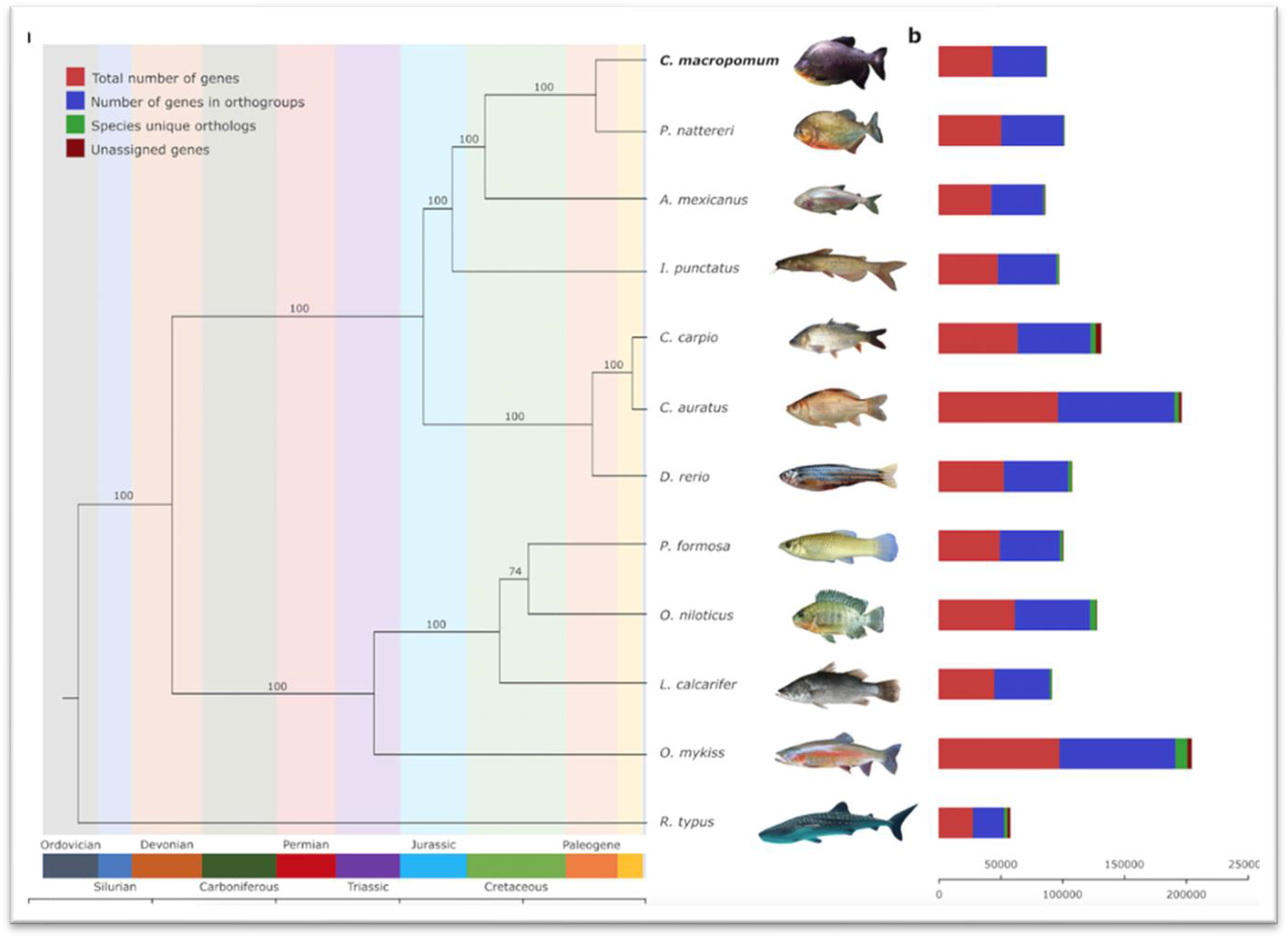
Whole-genome-predicted single copy orthologs phylogeny of 12 fish species including the newly sequenced genome of *C. macropomum.* To the right of the phylogeny bars show the proportion of different types of orthologs assigned by Orthofinder in each species.

## RE-USE POTENTIAL

Seasonal and long-term modifications in environmental conditions are well-acknowledged with periodic events of low water dissolved oxygen leading to hypoxia and even anoxia. Tambaqui is one of the amazon fish species that developed adaptions to deal with this, such as enlarging the lower lip to grasp oxygen better to survive in hypoxia. These, along with other fish adaptations to the Amazon aquatic ecosystem, are intriguing scientific questions that could be scientifically addressed using the present well-assembled and annotated tambaqui genome. Moreover, the availability of this annotated genome will pave the way to spur the development of tools for the genomic breeding programs of tambaqui, the most important native species for aquaculture in South America.

## AVAILABILITY OF SUPPORTING DATA

The datasets generated and analyzed during the current study are available on NCBI under the SRA numbers SRX10122091 to SRX10122101. The assembled genome and sequence annotations are on NCBI under the accession number GCF_904425465.1. The genome sequence and the annotation files - including CDS and proteins - can be downloaded from the NCBI FTP server (https://ftp.ncbi.nlm.nih.gov/genomes/all/GCF/904/425/465/GCF_904425465.1_Colossoma_macropomum/). A DataViewer can be accessed at https://www.ncbi.nlm.nih.gov/genome/gdv/browser/genome/?id=GCF_904425465.1.

## COMPETING INTERESTS

The authors declare no competing interests.

## AUTHOR CONTRIBUTIONS

AWSH, LLC, and DP designed and conceived this work; AWSH collected the samples; AWSH, MUS, DP, LLC, VMDAV wrote the manuscript; MUS and HM coordinated and carries out the bioinformatics analyses; AWSH, LLC and DP revised the manuscript. All authors read and approved the final manuscript.

## ACKNOWLEDGEMENTS

The authors acknowledge FAPESP (São Paulo Research Foundation # 2015/23883-0), National Council for Scientific and Technological Development (CNPq), and Coordination of Superior Level Staff Improvement (CAPES) through Tambaqui Project (Edital Pró-Amazonia – 047/2012) for financial support. CEPTA-ICMBio (Centro Nacional de Pesquisa e Conservação da Biodiversidade Aquática Continental) for tambaqui individual contribution to this work. Also, Dr. Andrew Veale for the critical review and English editing. AWSH, LLC, VWDAV, and DP are recipients of CNPq productivity scholarships (304662/2017-8, 304353/2019-1, 306718/2019-7, and 312273/2017-7, respectively).

## ADDITIONAL INFORMATION

Correspondence and requests for materials should be addressed to AWSH, DP or MUS.

## Notes

https://ftp.ncbi.nlm.nih.gov/genomes/all/GCF/904/425/465/GCF_904425465.1_Colossoma_macropomum/

### Competing Interest Statement

The authors have declared no competing interest.

