## Supplemental Figures and Table for "Genome assembly and annotation of the tambaqui (*Colossoma macropomum*): an emblematic fish of the Amazon River basin"

### Supplementary Material

**Figure S1.** BUSCO genes synteny of *Colosoma macropomum* (tambaqui; on the right side) and *Carassius auratus* (goldfish; on the left side). Synteny analysis of single copy genes reveal low conservation of homologous gene order between the species. The majority of *C. macropomum* genes are pulverized into several linkage groups of *C. auratus* genome, reflecting the different genome evolutionary events experienced by them.

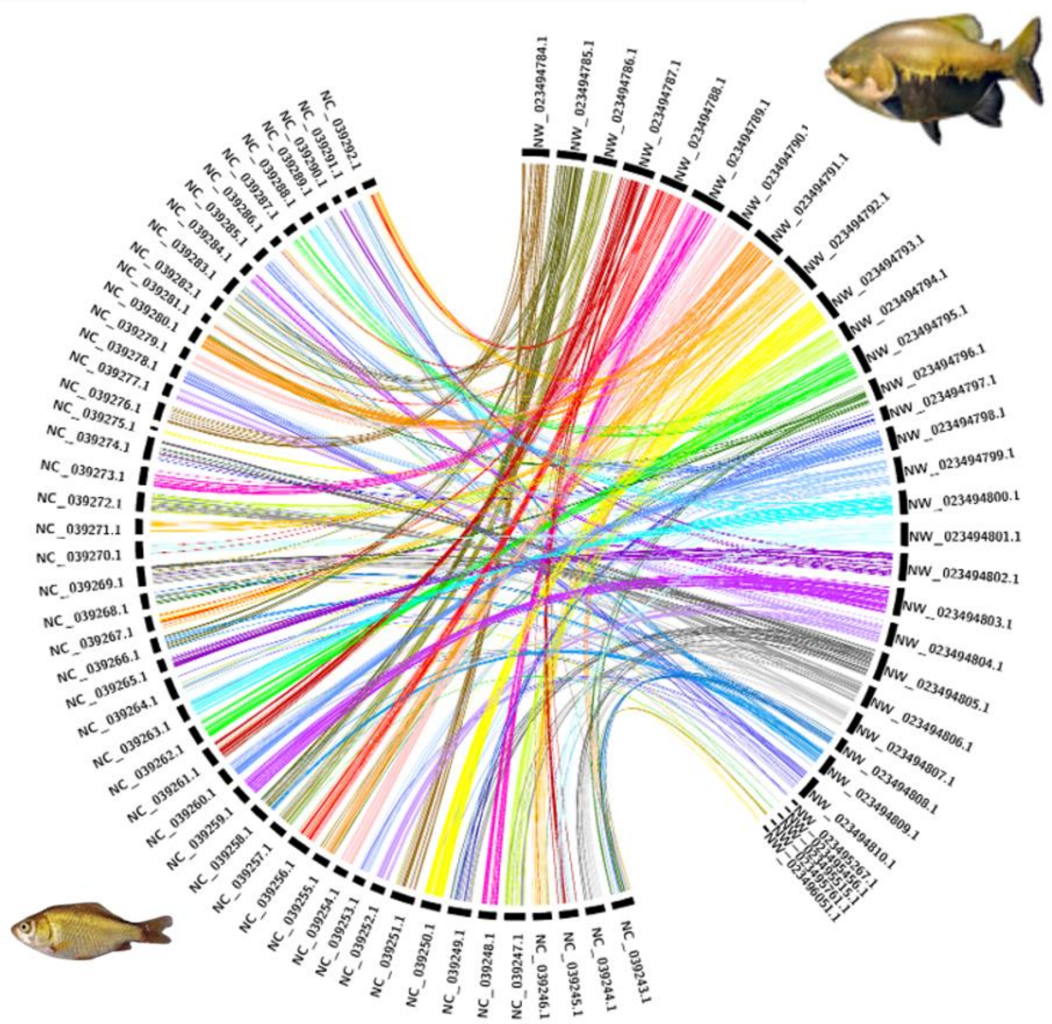

**Figure S2.** BUSCO genes synteny of *C. macropomum* (tambaqui; on the right side) and *Astyanax mexicanus* (Mexican tetra; on the left side). Synteny analysis confirm moderate conservation of homologous single copy genes order between these species. While large syntenic blocks persist, there is also a relatively large portion of *C. macropomum* genes which are fragmented into two or more linkage groups of *A. mexicanus* genome.

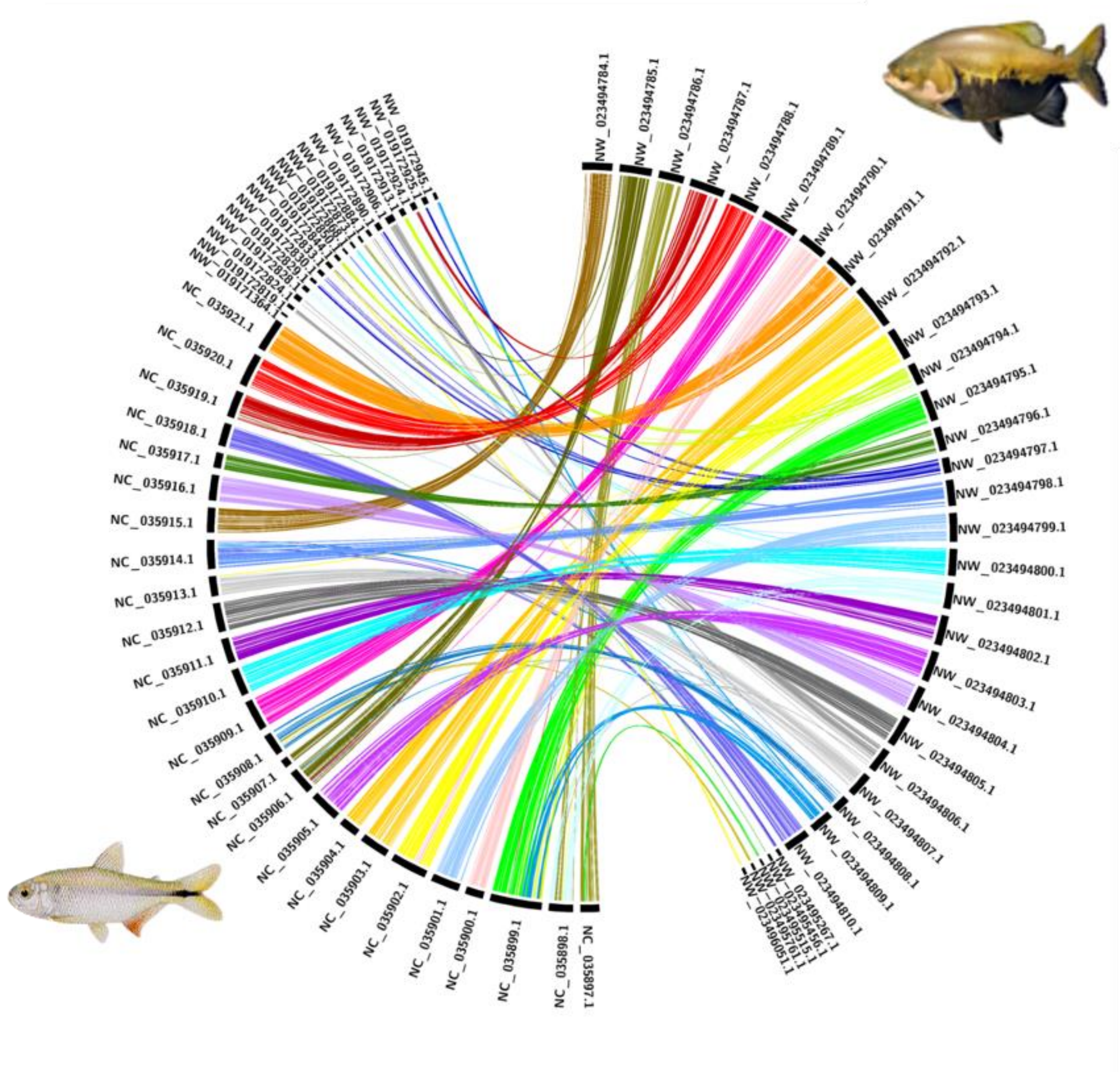

**Supplementary Information**

**Table S1**: NCBI accessions for the genes used for taxonomic ascertainment.

| Species | COI | TROP | fkh | RAG2 | sina |
|---|---|---|---|---|---|
| ***C. macropomum*** | HQ420845.1 | HQ420888.1 | AY817328.1 | AY804061.1 | AY790059.1 |
| ***P. brachypomus*** | HQ420838.1 | HQ420883.1 | AY817392.1 | AY804112.1 | AY790125.1 |
| ***P. mesopotamicus*** | HQ420837.1 | HQ420878.1 | AY817398.1 | AY804118.1 | AY790131.1 |
